# Consumption rate and dietary choice of cattle in species-rich mesic grasslands

**DOI:** 10.1101/2020.01.23.916635

**Authors:** Nóra Balogh, Béla Tóthmérész, Orsolya Valkó, Balázs Deák, Katalin Tóth, Zsolt Molnár, Csaba Vadász, Edina Tóth, Réka Kiss, Judit Sonkoly, Péter Török, Károly Antal, Júlia Tüdősné Budai, Tamás Miglécz, András Kelemen

## Abstract

For the improvement and maintenance of the desirable ecological value of grasslands it is necessary to manage them in a way which maintains their structure and their long-term functioning. Extensive grazing plays a crucial role in the seasonal biomass removal, thereby it prevents litter accumulation and shrub encroachment. Defoliation and biomass removal are among the most important effects of grazing on the vegetation, while the sufficient quantity and quality of plant biomass is an important ecosystem service for animal husbandry. In order to maintain the long term functioning of pastures it is important to gather information about the amount of consumed biomass and the dietary choice of the grazers. Therefore, we studied the direct effects of grazing on species-rich meadow steppes in Central-Hungary and the underlying mechanisms of dietary choice of cattle using trait-based approach. We asked the following questions: (i) What are the direct effects of grazing on the main biomass fractions (litter, moss, forbs and graminoids)? (ii) Which traits distinguish the preferred and non-preferred vascular plant species? The studied pastures were divided into two adjacent units, which were managed differently in the study year: the grazed units were managed by grazing for three months before the sampling date, while the control units remained ungrazed until the sampling. We collected above-ground biomass samples, measured leaf traits and shoot nitrogen content of plants. The consumption of the litter and moss biomass was negligible, while the reduction of the live biomass of vascular plants was 65%. Grazing significantly decreased the flowering success of plants. Cattle consumed species characterized by high specific leaf area and high nitrogen content. Based on our results we emphasize that, in order to ensure the reproduction of most plant species in the long term, it is unfavourable to graze an area every year in the same period. Instead, it is recommended to use grazing in a mosaic spatial and temporal pattern. The livestock carrying capacity of an area and the long-term management of grasslands can be carefully planned based on biomass measurements and the nutritional value of plants, which is well indicated by some easily measurable plant properties such as specific leaf area and the nitrogen content of species.

## 1. Introduction

The management of grasslands has undergone dramatic changes over the last centuries worldwide, both intensification of agricultural utilization and abandonment of traditional management practices led to the decline of their conservation value (Deák et al., 2017; Valkó et al., 2018). Moreover, the former area of grasslands has been fragmented and their ecological functions have been deteriorated, which makes them vulnerable (Bakker and Berendse, 1999; Kelemen et al., 2013). Thus, for the improvement and maintenance of the desirable ecological value of the grasslands it is necessary to manage them in a way which maintains their structure and their long-term functioning (Kelemen et al., 2014; Török et al., 2014; Tälle et al., 2016, Godó et al., 2017). Former studies proved that grazing adapted to the grassland type can be a proper tool to fulfil this aim (Isselstein et al., 2005; Tälle et al., 2016, Tölgyesi et al., 2015). Maintaining the appropriate management of pastures is much more efficient both from a nature conservation and a financial point of view than the reconstruction of the degraded stands (Török et al., 2011; Valkó et al., 2016). Extensive grazing plays a crucial role in the seasonal removal of biomass and prevents litter accumulation and shrub encroachment (Elias and Tischew, 2016). Grazing animals influence vegetation via several mechanisms; they can create colonization gaps and disperse the propagules by endo- and epizoochory allowing the establishment of many plant species (Will and Tackenberg, 2008; Eichberg and Donath 2018). Defoliation and biomass removal are among the most important effects of grazing, while the sufficient quantity and quality of plant biomass is an important ecosystem service for animal husbandry. In order to maintain the functioning of pastures in the long run, it is important to fine-tune grazing regimes according to the consumed biomass and the preference of the grazing animals.

Selective grazing may change the species composition and the trait distribution of the vegetation; species with particular trait syndromes may become more abundant while others may be suppressed (Tóth et al., 2018; Török et al., 2016). The scientific methods of vegetation studies have changed over time; recently the trait-based approaches became favoured over the former species identity-based approaches (Navas et al., 2010; Raevel et al., 2012). The trait-based approach offers an opportunity for the broad generalization of the findings, enabling the global-scale comparisons of results from habitats with distinct species pools (Lepš et al., 2011; Westoby, 1998; Kelemen et al., 2016). Several studies have shown that cattle prefer the nutrient-rich species, since their strategy is to maximise their energy intake rate with the lowest possible energy investment (Illius et al., 1992; Rutter et al., 2004; Soder et al., 2007). For this reason, plant species favoured by cattle generally have a high specific leaf area (SLA), high nitrogen and phosphorus content, and are often large-sized species that are visible from afar (Coppock et al., 1986; Díaz et al., 2001). Non-preferred species are often characterized by high dry matter content, high carbon/nitrogen ratio, and they are often hairy or thorny species (Moretto and Distel, 1997, Palkova and Lepš, 2008).

In many cases the efficient management of high-diversity pastures is hampered by the lack of knowledge about the functioning of these habitats. To fill this knowledge gap it is crucial to reveal the biomass consumption rate and dietary choice of grazers in these habitats. For this purpose, we studied the following questions: (i) What are the direct effects of grazing on the main biomass fractions (litter, moss, forbs and graminoids)? (ii) Which traits distinguish the preferred and non-preferred vascular plant species?

## 2. Material and methods

### 2.1. Study area and sampling

Two meadow steppe sites were sampled in the central part of the Great Hungarian Plain (Central-Hungary), in the Kiskunság National Park (coordinates for the centre: 47°06′N, 19°16′E). The climate in the region is continental, the mean annual temperature is 10 °C and the mean annual precipitation is 520 mm (Vadász et al., 2016). There are huge continuous primary and ancient steppes which cover several thousand hectares in this region. Their soils are meadow soils characterized by high humus content. These meadow steppes usually originated from *Molinion* meadows and are dominated by *Molinia caerulea, Chrysopogon gryllus* and *Agrostis stolonifera* (Molnár et al., 2008; Vadász et al., 2016). This habitat type is characterized by a unique species pool and high diversity, harbouring more than ten orchid species (e.g. *Anacamptis pyramidalis*, *Ophrys oestrifera*, *Ophrys sphegodes, Orchis coriophora*) and several other plants protected in Hungary (e.g. *Centaurea scabiosa* subsp. *sadleriana*, *Gentiana pneumonanthe*, *Iris sibirica*, *I. spuria*, *Koeleria javorkae*, *Ophioglossum vulgatum*, *Schoenus nigricans* and *Veratrum album*). The study sites have been managed by grazing for decades, nowadays extensive beef cattle grazing is typical with moderate grazing intensity (0.4 animal unit/ha). Both study sites were divided into two adjacent units which were managed differently in the study year, giving an opportunity to use split-plot design. The grazed units were grazed for three months before the sampling while the control units remained ungrazed in the study year until the sampling. Within each unit 35 above-ground biomass samples were collected from 20 cm × 20 cm plots in late June 2015, at the peak of biomass production. After drying, all 140 samples (2 sites, grazed and control units in both sites, 35 samples per unit) were separated to litter, moss and living biomass of vascular plants, and the latter fraction was sorted to species. Dry weights were measured with 0.01 g accuracy. The numbers of flowering shoots were also counted for each species in the biomass samples.

### 2.2. Data analyses

#### Community level analyses

In the community-level analyses, we considered all the vascular plant species detected in the biomass samples. We used linear mixed-effect models (LMEs) for exploring the effect of grazing on the dependent variables, where the grazing unit was set as fixed factor and the studied sites as random factor. Dependent variables were species density (species number/0.04m^2^), amount of litter (g/m^2^), moss biomass (g/m^2^), living biomass of vascular plants (g/m^2^), living biomass of graminoids (g/m^2^), living biomass of forbs (g/m^2^), total number of flowering species (species/0.04m^2^), number of flowering graminoid species (species/0.04m^2^), number of flowering forb species (species/0.04m^2^), total number of flowering shoots (shoot/m^2^), flowering shoot number of graminoids (shoot/m^2^) and flowering shoot number of forbs (shoot/m^2^).

#### Trait-based analyses

In order to reliably estimate species level biomass loss, we considered the species that occurred in more than 10% of the samples in each studied units. Altogether 100 vascular species were detected in the samples from which 29 species complied with this criterion, adding up to 92.5% of the total living biomass of vascular plants. To compare the biomass of these 29 species in the grazed units and controls, we used parametric and non-parametric pairwise tests (i.e. t-test, Welch t-test or Mann-Whitney U test) depending on the results of the normality tests (Shapiro-Wilk test) and the F-tests of the equality of variances. We classified these 29 species into three preference categories based on their biomass loss showed by the results of the pairwise tests as follows: (i) non-preferred: no significant differences in their biomass between the grazed and control units, (ii) moderately preferred: significantly higher biomass in control units, but with biomass loss lower than the average reduction of the living biomass of vascular plants (65%), (iii) highly preferred: significantly higher biomass in control units, with biomass loss greater than the average reduction of the living biomass of vascular plants. For the leaf trait measurements, we collected three individuals of each species and measured the following traits: leaf dry matter content (LDMC; %), leaf area (LA; mm^2^) and specific leaf area (SLA; mm^2^/mg). Moreover, we measured the nitrogen content of shoots (N; m/m%). We assigned shoot hairiness (present/absent) and shoot height of the species based on Király et al. (2009).

We applied one-way ANOVA with Fisher LSD post-hoc test to reveal the differences in the trait values characteristic to the three preference categories. We performed Principal component analysis (PCA) to study the covariance amongst traits. We tested the correlation between the species traits using Pearson correlation. We used R software for statistical analyses (R Core Team, 2017).

## 3. Results

### Community level analyses

LMEs did not reveal significant differences in the species richness of the grazed and control units. However, there were some short-lived species which were found only in the control units (e.g. *Bromus hordeaceus*, *Cerastium dubium*, *C. vulgare*, *Capsella bursa-pastoris* and *Myosotis stricta*). The differences between the control and the grazed units in the amount of litter and moss biomass were non-significant (Table 1). The living biomass of vascular plants decreased by 65.2% in grazed units compared to the controls, the reduction was similar for graminoid (65.5%) and forb (64.2%) biomass (Figure 2). LMEs showed that these reductions were significant, and there were no significant site effects detected (Table 1). Grazing significantly decreased the number of flowering species, both for graminoids and forbs (Table 1, Figure 3). The number of flowering shoots was reduced by 85.5% in the grazed units compared to controls; the reduction was 85.6% and 84.7% for graminoids and forbs, respectively (Table 1, Figure 3). There were no significant site effects detected in the case of flowering species and shoot numbers.

**Table 1.**
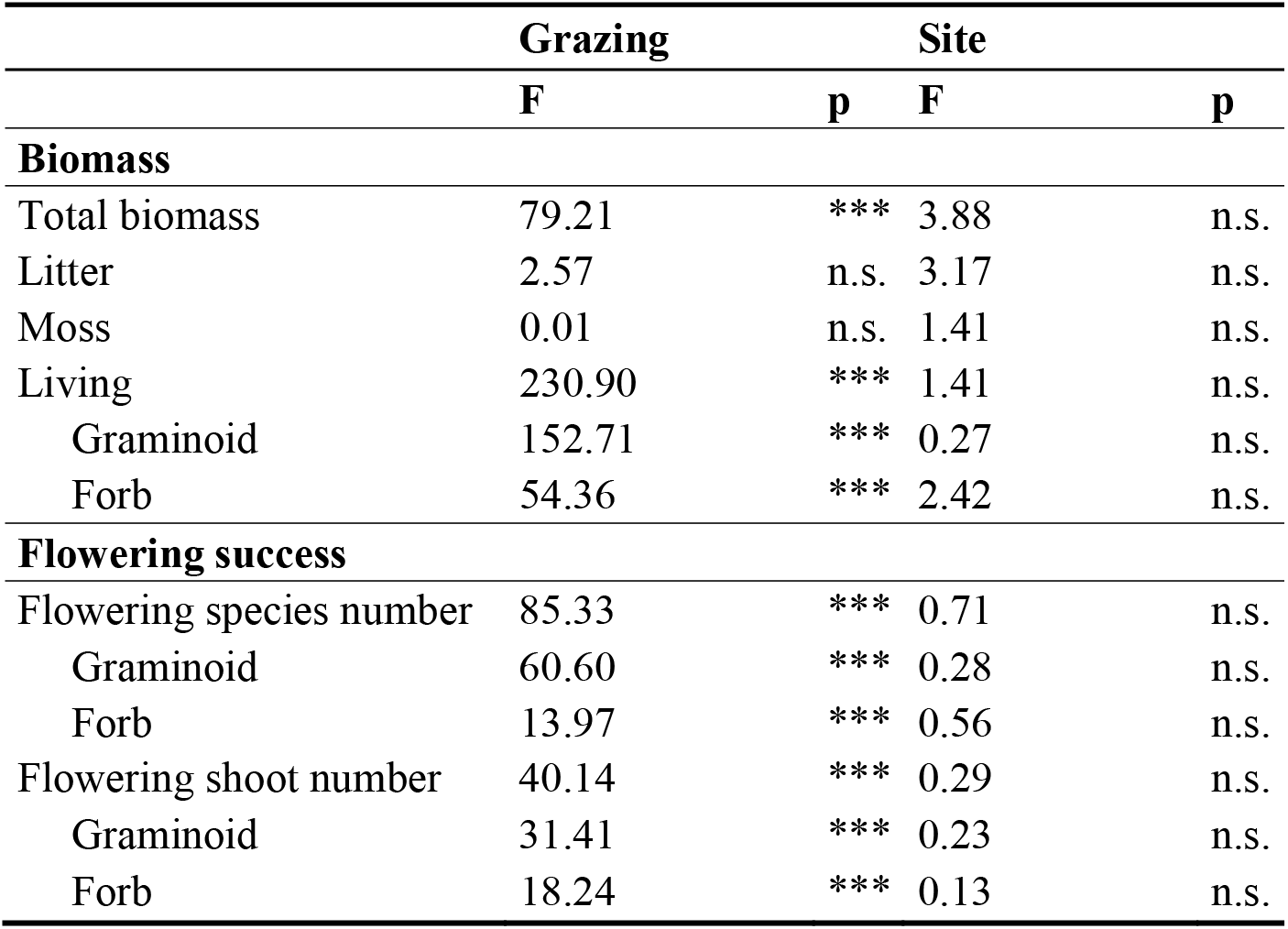
The effects of grazing on the biomass fractions and flowering success. The results of LMEs, where the grazing unit was set as fixed factor and the studied sites as random factor.

**Figure 1.**
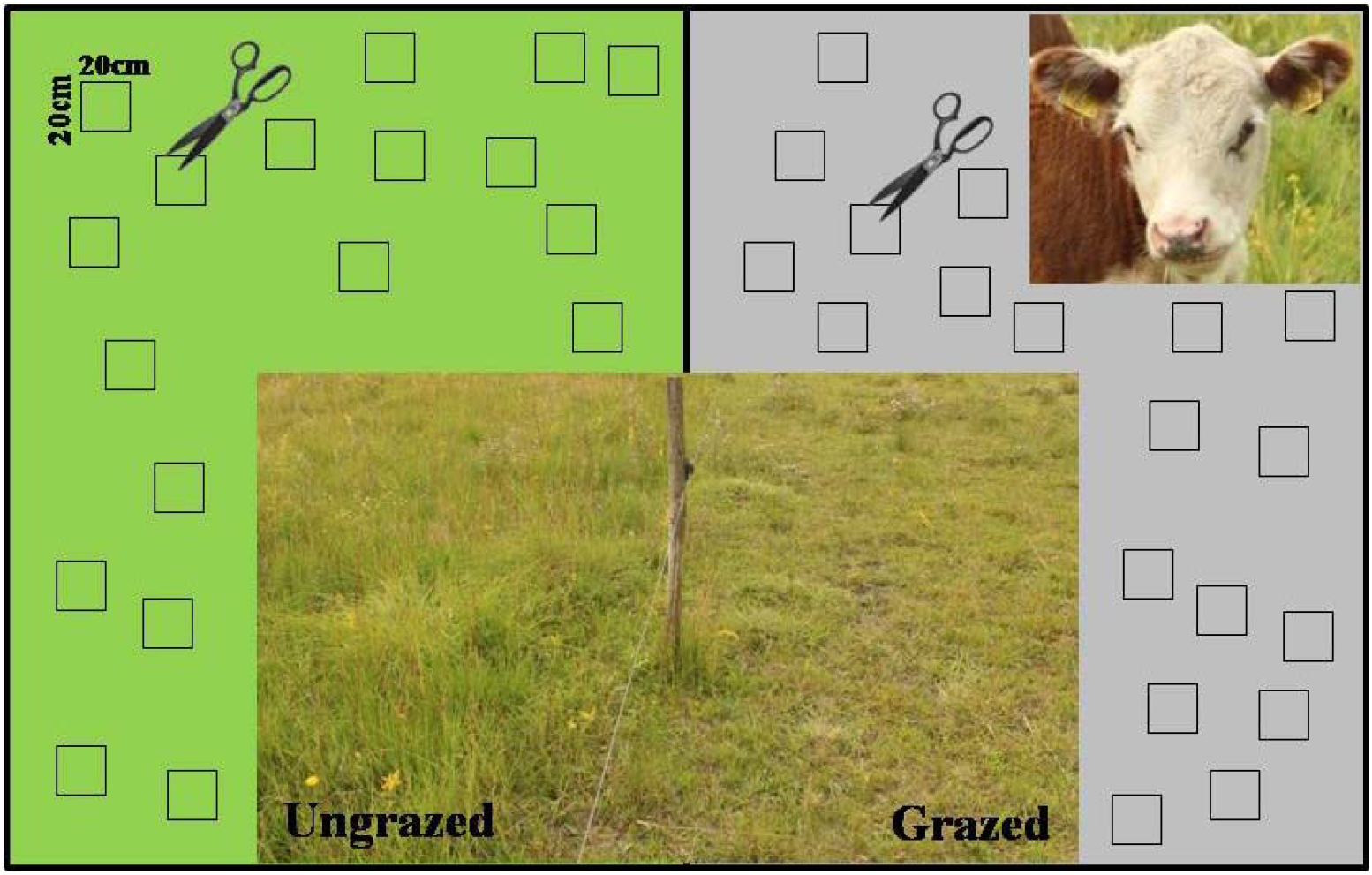
Our split-plot sampling design. The studied pastures were divided into two adjacent grazing units (i.e. grazed and ungrazed). Within each unit 35 above-ground biomass samples were collected from 20 cm × 20 cm plots.

**Figure 2.**
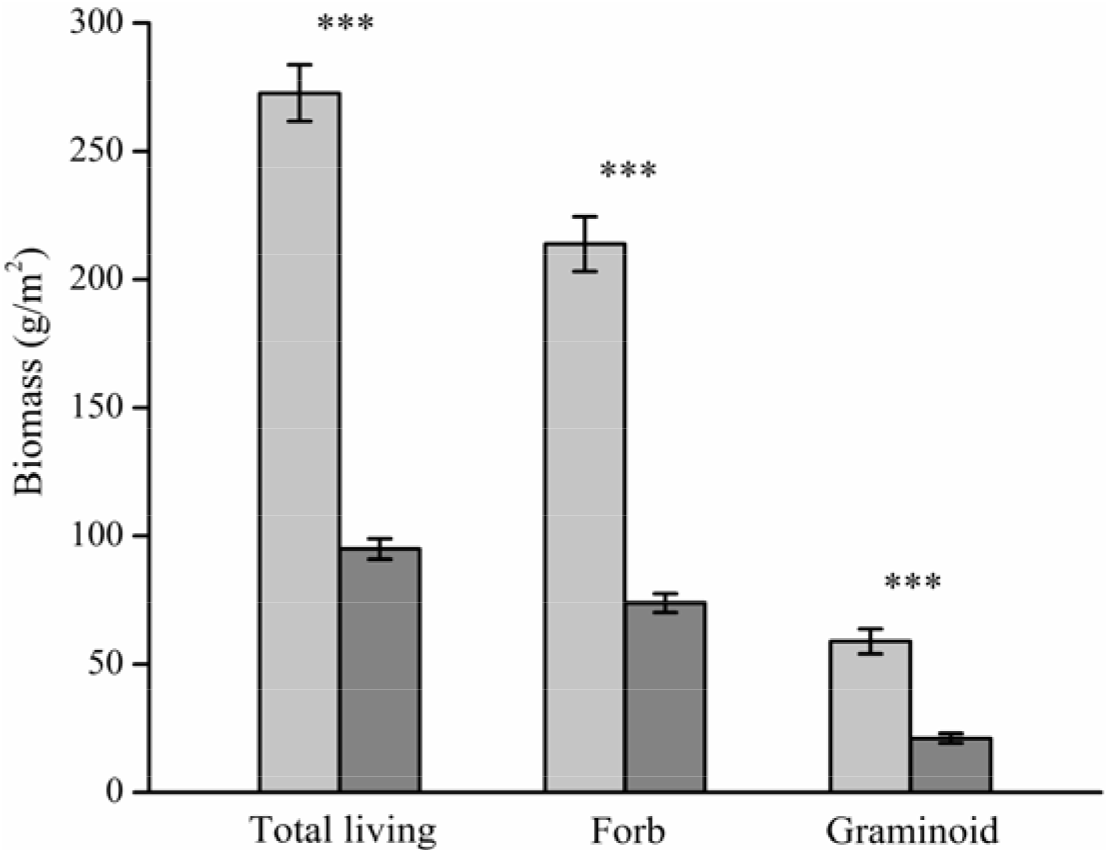
Living biomass fractions of ungrazed (control) and grazed units (mean ± SE). LMEs revealed significant grazing effects in the case of all living biomass fractions. Light bars: ungrazed units; dark bars: grazed units.

**Figure 3.**
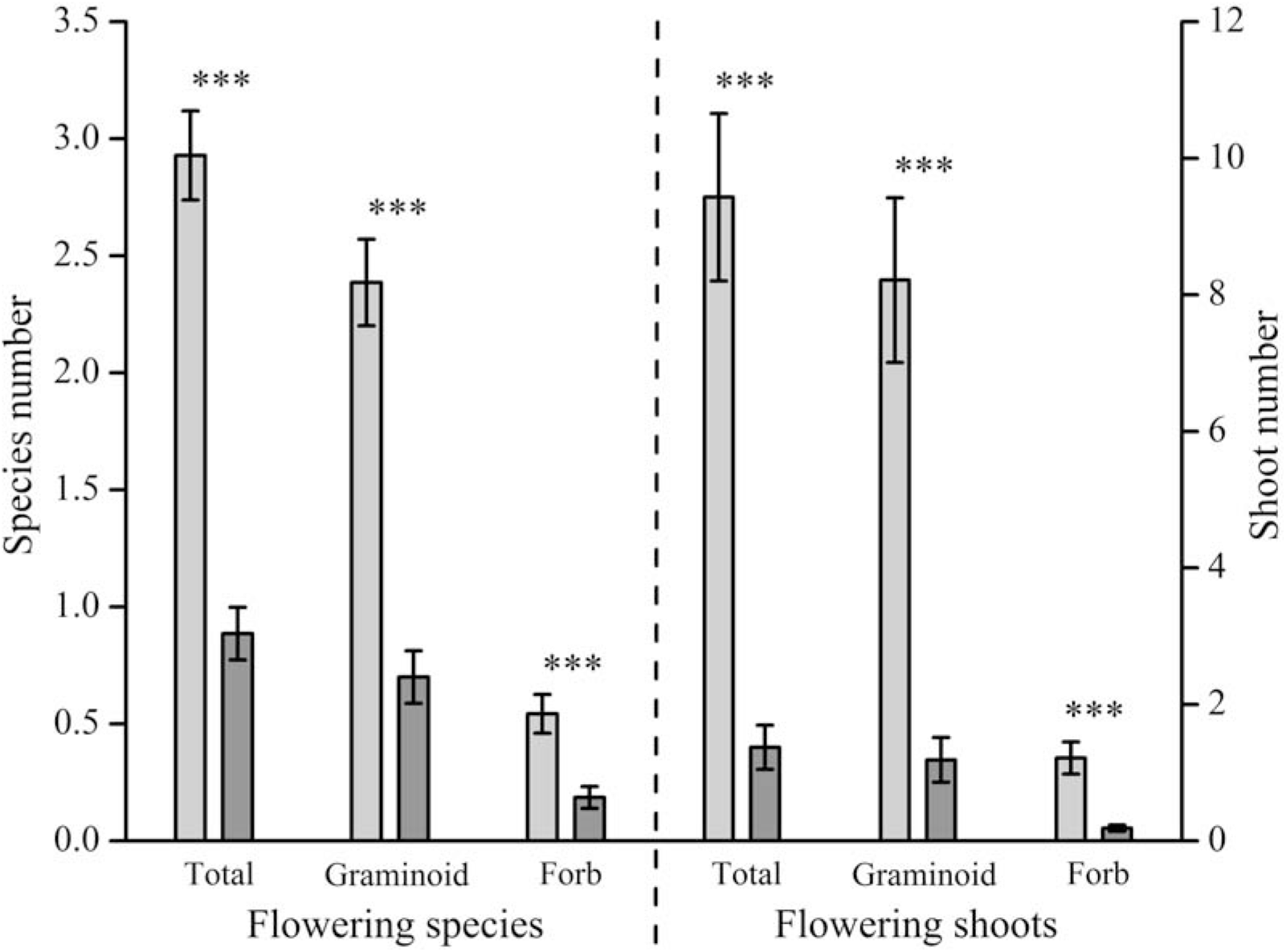
Differences in flowering success (No. of flowering species and No. of flowering shoots; mean ± SE) between the ungrazed (light bars) and grazed (dark bars) units. LMEs revealed significant grazing effects in all cases.

### Trait-based analyses

We classified the most frequent species into preference categories based on their biomass loss (Table 2) and compared the functional trait values of species belonging to different categories. The species of different categories significantly differed in their SLA (ANOVA: F=7.47; p<0.05) and nitrogen content (ANOVA: F=10.67; p<0.001); cattle preferred species with high SLA and nitrogen content (Fig. 4). There were no significant differences in other traits (LDMC, LA, plant height, hairiness) between the preference categories.

**Table 2.** Biomass reductions of the most frequent species. To compare their biomass between the grazed units and control, we used Welch t-test (W) or Mann-Whitney U test (M). We classified these species into three preference categories based on their biomass reduction and on the results of the pairwise tests. Notations: * - p < 0.05; ** - p <0.01; *** - p ≤ 0.001

| Species | Biomass reduction (%) | Test | Preference category |
| --- | --- | --- | --- |
| *Achillea asplenifolia* | 0.00 | M; n.s. | 1 |
| *Deschampsia caespitosa* | 0.00 | M; n.s. | 1 |
| *Festuca pseudovina* | 0.00 | M; n.s. | 1 |
| *Scirpoides holoschoenus* | 0.00 | W; n.s. | 1 |
| *Juncus articulatus* | 0.00 | M; n.s. | 1 |
| *Koeleria cristata* | 0.00 | M; n.s. | 1 |
| *Plantago lanceolata* | 0.00 | M; n.s. | 1 |
| *Plantago media* | 0.00 | M; n.s. | 1 |
| *Poa angustifolia* | 0.00 | M; n.s. | 1 |
| *Potentilla recta* | 0.00 | M; n.s. | 1 |
| *Schoenus nigricans* | 0.00 | M; n.s. | 1 |
| *Agrostis stolonifera* | 34.23 | M; * | 2 |
| *Carex flacca* | 64.83 | M; * | 2 |
| *Carex panicea* | 63.25 | W; ** | 2 |
| *Carex tomentosa* | 64.14 | W; * | 2 |
| *Chrysopogon gryllus* | 9.79 | M; * | 2 |
| *Dactylis glomerata* | 63.85 | W; ** | 2 |
| *Festuca pratensis* | 13.51 | M; ** | 2 |
| *Inula britannica* | 10.29 | M; ** | 2 |
| *Serratula tinctoria* | 56.07 | M; *** | 2 |
| *Elymus repens* | 76.98 | W; ** | 3 |
| *Centaurea jacea* | 79.47 | M; *** | 3 |
| *Daucus carota* | 95.41 | M; *** | 3 |
| *Galium verum* | 91.05 | M; *** | 3 |
| *Leontodon hispidus* | 87.21 | M; ** | 3 |
| *Lotus corniculatus* | 81.80 | M; * | 3 |
| *Molinia caerulea* | 92.57 | M; *** | 3 |
| *Picris hieracioides* | 77.41 | W; * | 3 |
| *Tetragonolobus maritimus* | 81.85 | M; * | 3 |

**Figure 4.**
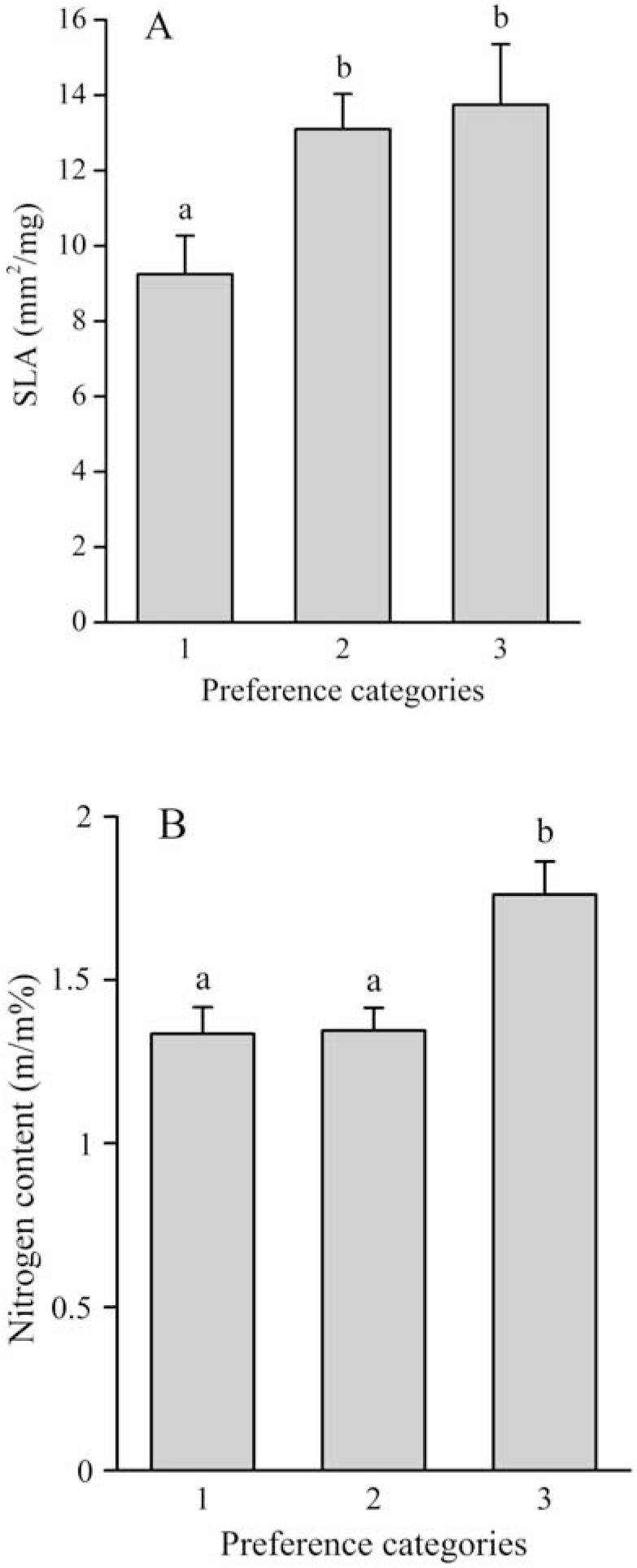
Differences in SLA and nitrogen content (means + SE) between the preference categories (1 – non-preferred; 2 – moderately preferred; 3 – highly preferred). Different lowercase letters denote significant differences revealed by one-way ANOVA and Fisher LSD post-hoc test (p < 0.05).

We revealed the association between the functional traits using PCA and Pearson correlations (Fig. 5). SLA was positively correlated with nitrogen content (Pearson: r= 0.47; p< 0.05) and negatively with LDMC (Pearson: r= −0.49; p< 0.01), and there was a negative correlation between LA and LDMC (Pearson: r= −0.46; p< 0.05).

**Figure 5.**
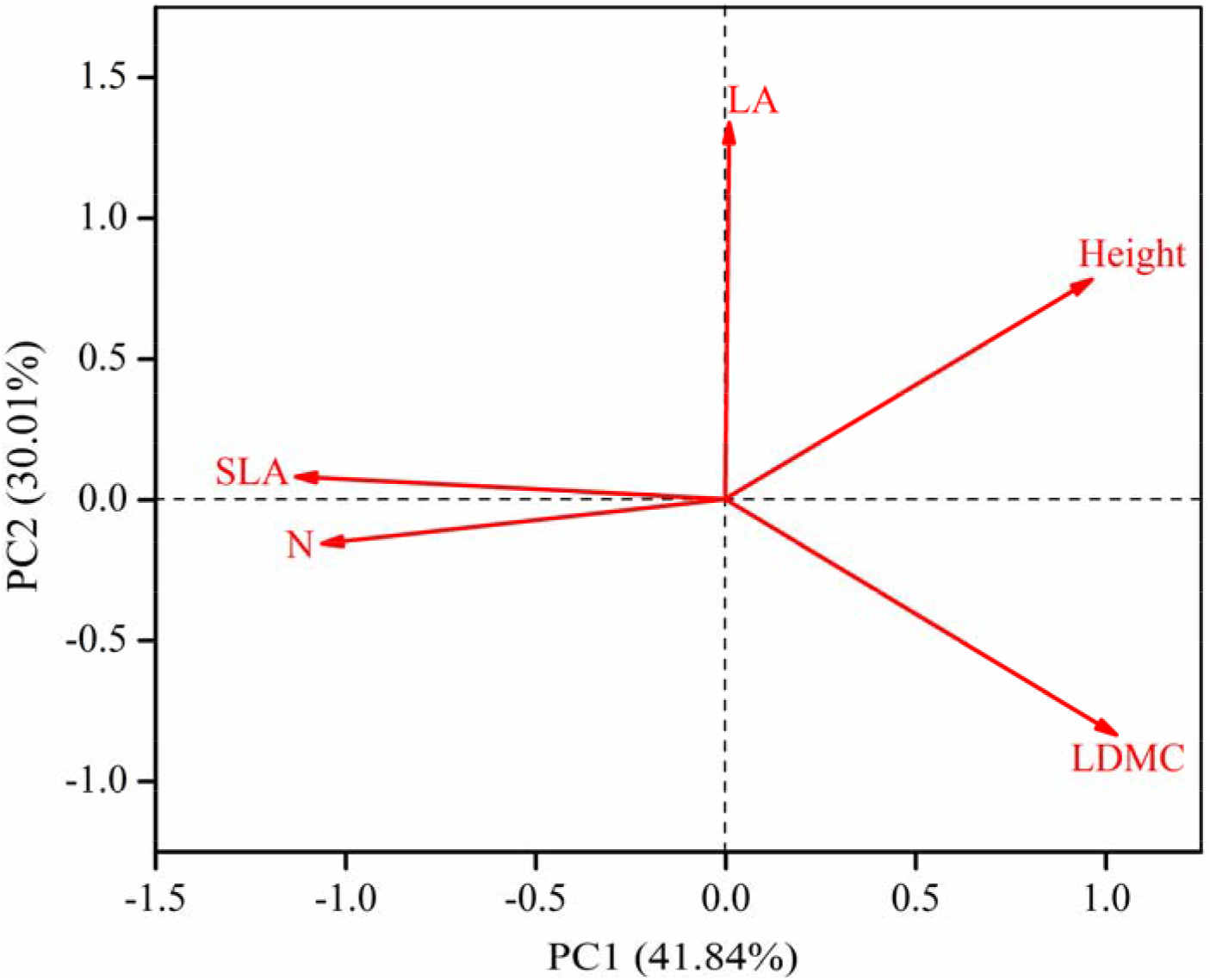
PCA ordination to present the covariance amongst the studied functional traits. Eigenvalues for 1st and 2nd axis: 2.09 and 1.50, respectively; explained variation: 41.84% and 30.01%, respectively. Abbreviations: LDMC – leaf dry matter content; LA – leaf area; SLA – specific leaf area; N – shoot nitrogen content; Height – shoot height.

## 4. Discussion

There are lot of studies about grazing effects on grasslands with contradictory results regarding its impact on species richness and biomass (Ferraro and Oesterheld, 2002; Matches, 1992). One of the main reasons for the various results is the different temporal scale of the studies. The direct effects of grazing (i.e. the consumption of plants), the short-term effects of grazing (i.e. the vegetation properties in the next year after grazing) and the long-term effects of grazing (i.e. changed species composition and vegetation structure) necessarily differ from each other (Asner et al., 2004). In this paper, we studied the direct effects of grazing, which can lead to a more detailed understanding of the rate of biomass consumption and the dietary choice strategies of cattle.

In this study we expected and found that the species richness scores did not differ in the grazed and control units, because the long-term management of the studied grazing units has been the same as they have been managed by grazing for decades. Therefore, we focused on the direct effects of grazing on the biomass and flowering success of plants.

In moderately grazed grasslands Plachter and Hampicke detected an approximately 45% decrease in litter amounts, while other studies typically found greater (70-85%) decreases in the long run (Kauffman et al., 2004; Bork et al., 2012). Our results correspond with the findings of Carilla et al. (2011) who demonstrated that not much litter is directly consumed by cattle. Our results also agree with the finding that the lower amount of litter observed on pastures is mainly due to the effects of the trampling and consumption of living biomass (Xiong and Nilsson, 1999; Welker et al., 2004).

The effect of grazing on mosses has been studied mainly in moss-dominated alpine vegetation or tundra (Memmot et al., 1998; Virtanen, 2000). Memmot et al. (1998) demonstrated that spring grazing reduced moss cover by 50% in temperate cold deserts with open vegetation. Based on our study, we can assume that in habitats where herbaceous vegetation is abundantly available for the cattle the consumption of mosses is negligible (Rupprecht et al., 2016). This is consistent with the results of Oldén and Halme (2016) as they found that bryophytes are generally not eaten by large herbivores.

Studies about the direct effects of moderate intensity grazing on vascular plants’ biomass have come to similar results to ours; grasslands showed a 60-70% reduction in living biomass due to the direct effect of grazing (Amiaud et al., 2008; Hofstede et al., 1995). However, in the literature, we can find examples for biomass consumption rates different from what we detected, for example Carilla et al. (2011) reported 4 to 10% consumption of the living biomass of vascular plants under moderate grazing intensity.

The knowledge on the productivity of grasslands and the dietary preferences of the livestock support the planning sustainable livestock farming. However, from the ecological viewpoint, we have to consider that the decrease of the flowering success due to grazing can be disproportionately higher than the rate of biomass loss as we detected in the present study. Although grazers often avoid consuming flowering individuals (particularly flowering graminoids), cattle repeatedly graze the majority of palatable plants before they reach the flowering state; thus, they decrease the efficiency of the generative propagation (Milchunas and Noy-Meir, 2002; Oesterheld and Oyarzábal, 2004; Mladek et al., 2013). Although the direct effect of grazing on the number of flowering shoots and flowering species is negative (Kimball and Schiffman, 2003), a distinction must be drawn between this direct effect (consumption of first grass) and even the short-term effects (e.g. flowering of regrowths or flowering in next year) (Plachter and Hampicke, 2010). Anderson and Frank (2003) reported that in regularly grazed areas, which have not yet been grazed in the given year, the number of flowering shoots is twice as high as in abandoned areas. A likely explanation of this phenomenon is that in grazed grasslands light penetration is usually higher than in non-grazed ones (Bakker et al., 2003), which is favourable for flowering. Therefore, the biomass allocated to the reproductive parts of plants relative to their vegetative parts is higher in grazed habitats (Niu et al., 2009). Based on the above results, we emphasize that, in order to ensure the reproduction of most species in the long term, it is unfavourable to graze an area every year in the same period. Instead, it is recommended to use a grazing regime which is mosaic in space and time (Vadász et al., 2016).

For the implementation of the mosaic management we have to consider the dietary choice and preference of the cattle. The differences in cattle’s preference can cause an uneven reduction in the biomass of different species, which can lead to a shift in the competitive interactions among species (Grant et al., 1996; Johnson and Sandercock, 2010). Different functional traits of species affect the vegetation composition via these community-level mechanisms. Grazers prefer highly digestible species with high nutritional value (Mladek et al., 2013; Moretto and Distel, 1997). These hard traits can be analysed via some soft traits as they positively correlate with SLA and nitrogen content (Mladek et al., 2013; Bullock et al., 2001). Our results justified this assumption as cattle preferred plants with high SLA and nitrogen content. These two traits were the only significant indicators of dietary preference.

The livestock carrying capacity of an area and the long-term management of grasslands can be carefully planned based on biomass measurements and the nutritional value of plants, which is well indicated by some easily measurable plant properties such as specific leaf area and nitrogen content. Moreover, it would be necessary to measure the amount of biomass in studies focusing on grazing effects, because using this method proper grazing intensities could be effectively determined for any given grassland.

## Acknowledgements

The authors were supported by MTA’s Post Doctoral Research Program (AK), Hungarian Scientific Research Fund (OV: NKFI FK 124404, NKFI KH 126476; BD: OTKA PD 115627; NKFI KH 130338, TM: NKFI PD 124548; PT: NKFIH K 119225; BT: OTKA K 116239, NKFI KH 126477, KT: NKFI PD 128302) and the Bolyai János Fellowship of the Hungarian Academy of Sciences (AK, OV, BD). We thank for the Kiskunság National Park Directorate for their support.

## References

1. Amiaud, B., Touzard, B., Bonis, A., Bouzillé, J.B., 2008. After grazing exclusion, is there any modification of strategy for two guerrilla species: *Elymus repens* (L.) Gould and *Agrostis stolonifera* (L.)? Plant Ecol. 197(1), 107–117. https://doi.org/10.1007/s11258-007-9364-z

2. Anderson, M.T., Frank, D.A., 2003. Defoliation effects on reproductive biomass: importance of scale and timing. Rangeland Ecol. Manage. 56(5), 501–516. https://doi.org/10.2458/azu_jrm_v56i5_anderson

3. Asner, G.P., Elmore, A.J., Olander, L.P., Martin, R.E., Harris, A.T., 2004. Grazing systems, ecosystem responses, and global change. Annu. Rev. Environ. Resour. 29, 261–299. https://doi.org/10.1146/annurev.energy.29.062403.102142

4. Bakker, J.P., Berendse, F., 1999. Constraints in the restoration of ecological diversity in grassland and heathland communities. Trends Ecol. Evol. 14(2), 63–68. https://doi.org/10.1016/S0169-5347(98)01544-4

5. Bakker, C., Blair, J.M., Knapp, A.K., 2003. Does resource availability, resource heterogeneity or species turnover mediate changes in plant species richness in grazed grasslands? Oecologia 137(3), 385–391. https://doi.org/10.1007/s00442-003-1360-y

6. Bork, E., Willms, W., Tannas, S., Alexander, M., 2012. Seasonal patterns of forage availability in the fescue grasslands under contrasting grazing histories. Rangeland Ecol. Manage. 65(1), 47–55. https://doi.org/10.2111/REM-D-11-00087.1

7. Bullock, J.M., Franklin, J., Stevenson, M.J., Silvertown, J., Coulson, S.J., Gregory, S.J., Tofts, R., 2001. A plant trait analysis of response to grazing in a long-tem experiment. J. Appl. Ecol. 38, 253–267. https://doi.org/10.1046/j.1365-2664.2001.00599.x

8. Carilla, J., Aragón, R., Gurvich, D.E., 2011. Fire and grazing differentially affect aerial biomass and species composition in Andean grasslands. Acta Oecol. 37(4), 337–345. https://doi.org/10.1016/j.actao.2011.03.006

9. Coppock, D.L., Swift, D.M., Ellis, J.E., 1986. Seasonal nutritional characteristics of livestock diets in a nomadic pastoral ecosystem. J. Appl. Ecol. 23, 585–595. https://doi.org/10.2307/2404038

10. Deák, B., Tölgyesi, Cs., Kelemen, A., Bátori, Z., Gallé, R., Bragina, T.M., Abil, Y.A., Valkó, O., 2017. The effects of micro-habitats and grazing intensity on the vegetation of burial mounds in the Kazakh steppes. Plant Ecol. Divers. 10, 509–520. https://doi.org/10.1080/17550874.2018.1430871

11. Díaz, S., Noy-Meir, I., Cabido, M., 2001. Can grazing response of herbaceous plants be predicted from simple vegetative traits? J. Appl. Ecol. 38, 497–508. https://doi.org/10.1046/j.1365-2664.2001.00635.x

12. Eichberg, C., Donath, T.W., 2018. Sheep trampling on surface◻lying seeds improves seedling recruitment in open sand ecosystems. Restor. Ecol. 26, S211–S219. https://doi.org/10.1111/rec.12650

13. Elias, D., Tischew, S., 2016. Goat pasturing – A biological solution to counteract shrub encroachment on abandoned dry grasslands in Central Europe? Agric. Ecosys. Environ. 234, 98–106. https://doi.org/10.1016/j.agee.2016.02.023

14. Ferraro, D.O., Oesterheld, M., 2002. Effect of defoliation on grass growth. A quantitative review. Oikos 98(1), 125–133. https://doi.org/10.1034/j.1600-0706.2002.980113.x

15. Godó, L., Valkó, O., Tóthmérész, B., Török, P., Kelemen, A., Deák, B., 2017. Scale-dependent effects of grazing on the species richness of alkaline and sand grasslands. Tuexenia 37, 229–246. https://doi.org/10.14471/2017.37.016

16. Grant, S.A., Torvell, L., Sim, E.M., Small, J.L., Armstrong, R.H., 1996. Controlled grazing studies on *Nardus* grassland: effects of between-tussock sward height and species of grazer on *Nardus* utilization and floristic composition in two fields in Scotland. J. Appl. Ecol., 1053–1064. https://doi.org/10.2307/2404685

17. Hofstede, R.G., Rossenaar, A.J., 1995. Biomass of grazed, burned, and undisturbed paramo grasslands, Colombia. II. Root mass and aboveground: belowground ratio. Arctic Alpine Res. 27(1), 13–18. https://doi.org/10.1080/00040851.1995.12003092

18. Illius, A.W., Clark, D.A., Hodgson, J., 1992. Discrimination and patch choice by sheep grazing grass-clover swards. J. Anim. Ecol. 61, 183–194. https://doi.org/10.2307/5521

19. Isselstein, J., Jeangros, B., Pavlu, V., 2005. Agronomic aspects of biodiversity targeted management of temperate grasslands in Europe – a review. Agronomy Res. 3(2), 139–151.

20. Johnson, T.N., Sandercock, B.K., 2010. Restoring tallgrass prairie and grassland bird populations in tall fescue pastures with winter grazing. Rangeland Ecol. Manage. 63(6), 679–688. https://doi.org/10.2111/REM-D-09-00076.1

21. Kauffman, J.B., Thorpe, A.S. and Brookshire, E.J., 2004. Livestock exclusion and belowground ecosystem responses in riparian meadows of eastern Oregon. Ecol. Appl. 14(6), 1671–1679.

22. Kelemen, A., Török, P., Valkó, O., Miglécz, T., Tóthmérész, B., 2013. Mechanisms shaping plant biomass and species richness: plant strategies and litter effect in alkali and loess grasslands. J. Veg. Sci. 24, 1195–1203. https://doi.org/10.1111/jvs.12027

23. Kelemen, A., Török, P., Valkó, O., Deák, B., Miglécz, T., Tóth, K., Ölvedi, T., Tóthmérész, B., 2014. Sustaining recovered grasslands is not likely without proper management: vegetation changes and large-scale evidences after cessation of mowing. Biodivers. Conserv. 23, 741–751. https://doi.org/10.1007/s10531-014-0631-8

24. Kelemen, A., Valkó, O., Kröel-Dulay, Gy., Deák, B., Török, P., Tóth, K., Miglécz, T., Tóthmérész, B., 2016. The invasion of common milkweed (*Asclepias syriaca*) in sandy old-fields – Is it a threat to the native flora? Appl. Veg. Sci. 19, 218–224. https://doi.org/10.1111/avsc.12225

25. Kimball, S., Schiffman, P.M., 2003. Differing effects of cattle grazing on native and alien plants. Conserv. Biol. 17(6), 1681–1693. https://doi.org/10.1111/j.1523-1739.2003.00205.x

26. Király, G., 2009. Újmagyar Füvészkönyv. Magyarország hajtásos növényei. (New Hungarian Herbal. The vascular plants of Hungary. Identification key.). Aggtelek National Park Directorate, Jósvafő.

27. Lepš, J., de Bello, F., Šmilauer, P., Doležal, J., 2011. Community trait response to environment: disentangling species turnover vs intraspecific trait variability effects. Ecography 34(5), 856–863. https://doi.org/10.1111/j.1600-0587.2010.06904.x

28. Matches, A.G., 1992. Plant response to grazing: a review. J. Product. Agric. 5(1), 1–7. https://doi.org/10.2134/jpa1992.0001

29. Memmott, K.L., Anderson, V.J., Monsen, S.B., 1998. Seasonal grazing impact on cryptogamic crusts in a cold desert ecosystem. Rangeland Ecol. Manage. 51(5), 547–550.

30. Milchunas, D.G., Noy◻Meir, I., 2002. Grazing refuges, external avoidance of herbivory and plant diversity. Oikos 99(1), 113–130. https://doi.org/10.1034/j.1600-0706.2002.990112.x

31. Mladek, J., Mladkova, P., Hejcmanová, P., Dvorský, M., Pavlu, V., De Bello, F., Duchoslav, M., Hejcman, M., Pakeman, R.J., 2013. Plant trait assembly affects superiority of grazer’s foraging strategies in species-rich grasslands. PloS ONE 8(7), e69800. https://doi.org/10.1371/journal.pone.0069800

32. Molnár, Z., Biró, M., Bölöni, J., Horváth, F., 2008. Distribution of the (semi-)natural habitats in Hungary I. Marshes and grasslands. Acta Bot. Hung. 50, 59–105. https://doi.org/10.1556/ABot.50.2008.Suppl.5

33. Moretto, A.S., Distel, R.A., 1997. Competitive interactions between palatable and unpalatable grasses native to a temperate semi-arid grassland of Argentina. Plant Ecol. 130, 155–161. https://doi.org/10.1023/A:1009723009012

34. Navas, M.L., Roumet, C., Bellmann, A., Laurent, G., Garnier, E., 2010. Suites of plant traits in species from different stages of a Mediterranean secondary succession. Plant Biol. 12(1), 183–196. https://doi.org/10.1111/j.1438-8677.2009.00208.x

35. Niu, K., Choler, P., Zhao, B., Du, G., 2009. The allometry of reproductive biomass in response to land use in Tibetan alpine grasslands. Funct. Ecol. 23(2), 274–283. https://doi.org/10.1111/j.1365-2435.2008.01502.x

36. Oesterheld, M., Oyarzábal, M., 2004. Grass-to-grass protection from grazing in a semi-arid steppe. Facilitation, competition, and mass effect. Oikos 107(3), 576–582. https://doi.org/10.1111/j.0030-1299.2004.13442.x

37. Oldén, A., Halme, P., 2016. Microhabitat determines how grazing affects bryophytes in wood-pastures. Biodivers. Conserv. 25(6), 1151–1165. https://doi.org/10.1007/s10531-016-1115-9

38. Pálková, K., Lepš, J., 2008. Positive relationship between plant palatability and litter decomposition in meadow plants. Community Ecol. 9(1), 17–27. https://doi.org/10.1556/ComEc.9.2008.1.3

39. Plachter, H., Hampicke, U., (Eds.). 2010. Large-scale livestock grazing: a management tool for nature conservation. Springer Science & Business Media.

40. R Core Team. (2017). R: A Language and Environment for Statistical Computing. Vienna: R Foundation for Statistical Computing.

41. Raevel, V., Violle, C., Munoz, F., 2012. Mechanisms of ecological succession: insights from plant functional strategies. Oikos 121(11), 1761–1770. https://doi.org/10.1111/j.1600-0706.2012.20261.x

42. Rupprecht, D., Gilhaus, K., Hölzel, N. 2016. Effects of year-round grazing on the vegetation of nutrient-poor grass-and heathlands – Evidence from a large-scale survey. Agriculture, Ecosystems & Environment, 234, 16–22. https://doi.org/10.1016/j.agee.2016.02.015

43. Rutter, S.M., Orr, R.J., Yarrow, N.H., Champion, R.A., 2004. Dietary preference of dairy cows grazing ryegrass and white clover. J. Dairy Sci. 87(5), 1317–1324. https://doi.org/10.3168/jds.S0022-0302(04)73281-6

44. Soder, K.J., Rook, A.J., Sanderson, M.A., Goslee, S.C., 2007. Interaction of plant species diversity on grazing behavior and performance of livestock grazing temperate region pastures. Crop Sci. 47(1), 416–425. https://doi.org/10.2135/cropsci2006.01.0061

45. Tälle, M., Deák, B., Poschlod, P., Valkó, O., Westerberg, L., Milberg, P., 2016. Grazing vs. mowing: a meta-analysis of biodiversity benefits for grassland management. Agric. Ecosyst. Environ. 15, 200–212. https://doi.org/10.1016/j.agee.2016.02.008

46. Teuber, L.M., Hölzel, N., Fraser, L.H., 2013. Livestock grazing in intermountain depressional wetlands – Effects on plant strategies, soil characteristics and biomass. Agric. Ecosyst. Environ. 175, 21–28. https://doi.org/10.1016/j.agee.2013.04.017

47. Tölgyesi, C., Bátori, Z., Erdős, L., Gallé, R., Körmöczi, L., 2015. Plant diversity patterns of a Hungarian steppe-wetland mosaic in relation to grazing regime and land use history. Tuexenia 35, 399–416. https://doi.org/10.14471/2015.35.006

48. Török, P., Vida, E., Deák, B., Lengyel, Sz., Tóthmérész, B., 2011. Grassland restoration on former croplands in Europe: as assessment of applicability of techniques and costs. – Biodivers. Conserv. 20, 2311–2332. https://doi.org/10.1007/s10531-011-9992-4

49. Török, P., Valkó, O., Deák, B., Kelemen, A., Tóthmérész, B., 2014. Traditional cattle grazing in a mosaic alkali landscape: Effects on grassland biodiversity along a moisture gradient. – PLoS ONE 9(5), e97095. https://doi.org/10.1371/journal.pone.0097095

50. Török, P., Valkó, O., Deák, B., Kelemen, A., Tóth, E., Tóthmérész, B., 2016. Managing for composition or species diversity? – Pastoral and year-round grazing systems in alkali grasslands. Agric. Ecosyst. Environ. 234, 23–30. https://doi.org/10.1016/j.agee.2016.01.010

51. Tóth, E., Deák, B., Valkó, O., Kelemen, A., Miglécz, T., Tóthmérész, B., Török, P., 2018. Livestock type is more crucial than grazing intensity: Traditional cattle and sheep grazing in short-grass steppes. Land Degrad. Develop. 29(2), 231–239. https://doi.org/10.1002/ldr.2514

52. Vadász, C., Máté, A., Kun, R., Vadász-Besnyői, V., 2016. Quantifying the diversifying potential of conservation management systems: An evidence-based conceptual model for managing species-rich grasslands. Agric. Ecosyst. Environ. 234, 134–141. https://doi.org/10.1016/j.agee.2016.03.044

53. Valkó, O., Deák, B., Török, P., Kelemen, A., Miglécz, T., Tóth, K., Tóthmérész, B., 2016. Abandonment of croplands: problem or chance for grassland restoration? Case studies from Hungary. Ecosys. Health Sustain. 2(2), e01208. https://doi.org/10.1002/ehs2.1208

54. Valkó, O., Venn, S., Zmihorski, M., Biurrun, I., Labadessa, R., Loos, J., 2018. The challenge of abandonment for the sustainable management of Palaearctic natural and semi-natural grasslands. Hacquetia 17(1), 5–16. https://doi.org/10.1515/hacq-2017-0019

55. Virtanen, R., 2000. Effects of grazing on above◻ground biomass on a mountain snowbed, NW Finland. Oikos 90(2), 295–300. https://doi.org/10.1034/j.1600-0706.2000.900209.x

56. Welker, J.M., Fahnestock, J.T., Povirk, K.L., Bilbrough, C.J., Piper, R.E., 2004. Alpine grassland CO2 exchange and nitrogen cycling: grazing history effects, Medicine Bow Range, Wyoming, USA. Arct. Antarct. Alp. Res. 36(1), 11–20. https://doi.org/10.1657/1523-0430(2004)036[0011:AGCEAN]2.0.CO;2

57. Westoby, M., 1998. A leaf-height-seed (LHS) plant ecology strategy scheme. Plant Soil 199(2), 213–227. https://doi.org/10.1023/A:1004327224729

58. Will, H., Tackenberg, O., 2008. A mechanistic simulation model of seed dispersal by animals. J. Ecol. 96(5), 1011–1022. https://doi.org/10.1111/j.1365-2745.2007.01341.x

59. Xiong, S., Nilsson, C., 1999. The effects of plant litter on vegetation: a metaanalysis. J. Ecol. 87(6), 984–994. https://doi.org/10.1046/j.1365-2745.1999.00414.x

